# First Attempt to Identify and Map QTLs Associated with Promiscuous Nodulation in Soybean

**DOI:** 10.1101/688028

**Authors:** Eric E. Agoyi, John B. Tumuhairwe, Godfree Chigeza, Phinehas Tukamuhabwa, Brian W. Diers

## Abstract

To inform possibility of conducting marker assisted breeding of promiscuous soybean varieties, this study used 92 F2 lines from biparental cross to identify QTLs associated with promiscuous nodulation in soybean. GBS; genotyping by sequencing platform was used to generate SNPs through the pipeline 2 in TASSEL 5.0, Bowtie2 version 2.2.8 for tag alignment, Beagle version 4.1 to impute missing SNPs and R-QTL package in R for QTL identification. Four nodulation traits were assessed ***viz*** number of nodules (NN), percent of effective nodules (NE), fresh weight and dry weight of nodules (NFW and NDW). Two QTLs were identified on chromosomes 10 and 13. Both QTLs were associated with NN, only QTL13 was associated NE and only QTL10 was associated with nodule ‘weights. It was observed that NN, NFW and NDW shared QTL10 and NN and NE share QTL13 allowing hypothetize on the existence of pleiotropic genes in those those two regions. Over dominance effect was observed for QTL10 and non additive effect for QTL13. The paper recommend investigations be pursued to validate those QTLs and set foundation for marker assisted selection of promiscuous soybean varieties. Also these findings could serve as starting point for gene cloning to better understand nodulation trait in soybean.

## INTRODUCTION

Soybean (***Glycine max*** L. Merrill) is a grain legume with high protein content making it an alternative protein source in diet formulation, globally and particularly in Africa where there is low affordability of animal proteins. The crop requires high doses of nitrogen to process these proteins mostly at pod filling stage (Machido ***et al***., 2011). It has been reported that soybean crops require up to 80 kg of nitrogen per hectare to yield a ton of grain, with approximatively 58% of this nitrogen coming from N_2_ fixation (Salvagotti ***et al***., 2008), which is estimated up to 337 kg of nitrogen per hectare when inoculated with effective strains of ***Bradyrhizobium***. Incidentally, soybean is a highly specific legume making symbiotic relationship mainly with ***Bradyrhizobia japonicum***. This specific symbiotic association reduces on the nitrogen fixation potential of most soybean cultivars, especially in areas where ***Bradyrhizobia japonicum*** is not endemic. Breeding for promiscuity in soybean is prioritized to increase biological nitrogen fixation and improve production (IITA, 1996).

Like most economically important traits, promiscuous nodulation in soybean is likely to be quantitative, evidence of this has been shown by Agoyi ***et al.*** (2016b) who reported predominant additive gene actions in promiscuous nodulation traits in soybean. Quantitative traits are controlled by genes located in regions known as quantitative trait loci (QTLs) (Prasanna, 2007). In dealing with quantitative traits, molecular breeding requires the mapping of QTLs associated with the traits under consideration, to enable marker assisted breeding and individual gene cloning. QTL mapping is the process of constructing a linkage map and analyzing QTLs with the aim of identifying their number, where there are located, and their effect on traits of interest (Doerge, 2002; Jonah ***et al.***, 2011). QTLs analysis aims at identifying molecular markers that are closely linked to the desired allele, it is useful for marker-assisted breeding, as it provides knowledge for detecting marker-trait association (Jiang, 2013). According to Malosetti ***et al***. (2011) mapping QTLs helps in dissecting the complexity of quantitative traits by identifying loci with effect on the trait. It is therefore a major step towards cloning individual genes related to economically important traits. This has not been exploited on promiscuous nodulation in soybean.

Population types used for QTLs mapping include F2_s_, Backcrosses, recombinant inbred lines (RILs), and near-isogenic lines (NILs). Parental lines used to generate any of those populations must be genetically distant to show polymorphism (Semagn ***et al***., 2006).

An appropriate and cheap high-throughput genotyping platform such as next generation sequencing (NGS) is key for efficient mapping of QTLs and genomic studies. In this study we used Genotyping by sequencing (GBS), an ideal platform that shows efficiency from single gene markers to whole genome profiling (Poland and Rife, 2012). Several authors have used GBS to serve various purposes; it has been used for genome sequencing and gene mapping, genomic diversity studies, genome wide association studies (GWAS), genomic selection (GS), or improve marker resolution in number of crops including soybean (Yu ***et al.***, 2015; Rincker ***et al.***, 2016), maize (Romay ***et al***., 2013), wheat (van Poecke ***et al***., 2013), barley (Fu and Peterson, 2011), rice, potato (Uitdewilligen ***et al***., 2013), switch grass (Lu ***et al***., 2013), yellow mustard (Fu ***et al***., 2014) and Brassica oleracea (Liu ***et al***., 2014). GBS uses a multiplexed system for constructing reduced representation libraries for high diversity and large genome species (Elshire ***et al***., 2011). GBS generates hundreds of millions of single nucleotide polymosphisms (SNPs) for use in genetic analyses and genotyping (Beissinger ***et al***., 2013). It is well suited for dealing with reduced sample size, as barcoded DNA from the different samples are pooled. GBS requires fewer PCR and purification steps, barcoding is efficient and easy to operate, with no limitation due to lack of reference genome (Davey ***et al***., 2011). GBS allows SNPs to be discovered and genotyped at the same time (Poland and Rife, 2012; Narum ***et al***., 2013). GBS has enabled identifying a set of 205,614 SNPs in soybean, thus providing valuable tools for soybean breeding programs (Lam ***et al***., 2010).

The present study identified and mapped QTLs associated to promiscuous nodulation in soybean, to provide basis for marker-assisted selection and nodulation genes cloning.

## MATERIAL AND METHODS

### MAPPING POPULATION

The parents used consisted of two soybean genotypes Wondersoya and GC 2043 that are genetically distant in their response to ***Bradyrhizobium sp.*** strain USDA 3456. For instance, Wondersoya is highly responsive to ***B***. sp USDA 3456, while GC 2043 showed low response to that ***Bradyrhizobium*** strain (Agoyi ***et al***., 2016a). ***Bradyrhizobium*** spp which have been reported as native and widespread in African soils (Abaidoo ***et al***., 2000) are known to form nodules with promiscuous soybean genotypes (IITA, 1996).

A reciprocal cross was made between Wondersoya and GC 2043, F1 seeds were grown together with their corresponding female parent in plastic pots to generate F2 seeds and at the same time to check the successfulness of the crosses. Morphological traits such as hypocotyl color, pubescence color on stem, leaf and pod, days to flowering, flower color, maturity stage, plant height, were used to eliminate selfed individuals. Later on, seeds from parental lines and F2 seeds were inoculated with fresh culture of ***Bradyrhizobium sp.*** strain USDA 3456. ***Bradyrhizobium sp.*** strain USDA 3456 was obtained from Biofix (Kenya), purified and incubated in Soil Science Biological Nitrogen Fixation (BNF) laboratory at Makerere University. It was grown to 7.91 × 109 cells g^−1^ and formulated into an inoculum carried in steam-sterilised peat soil. Two table spoonfuls of sugar were dissolved into 300 ml of clean lukewarm water, in a soda bottle, to be used as sticker. The inoculant was mixed with the sticker and directly applied on seeds to enhance plant-rhizobium association. Inoculated seeds were planted in plastic pots filled up with steam-sterilized top soil in screenhouse at Makerere University Agricultural Research Institute of Kabanyolo (MUARIK). Prior to steam sterilization, soil samples were analyzed for nutrients content. As it was established in earlier studies that P and K are essential for biological nitrogen fixation (Giller et al., 1997), the soil was pre-mixed with 0.0356 and 0.036 g of TSP (Triple Super Phosphate) and Muriate of Potash (MOP), respectively per kg of soil, representing 20 kg of phosphorus and 40 kg of potassium per hectare. Ninety-two individual plants were successfully generated.

### Phenotyping F2 population

Six weeks after emergence, each of the 92 individual plants were evaluated for nodulation. Each plant was carefully released from the soil aggregate, root system was carefully and abundantly washed, rinsed and wrapped in tissue paper to reduce wetness. All nodules were harvested per plant and counted for number of nodules (NN), and weighed to determine their fresh weight (NFW). Then, all nodules were split opened using bistoury to assess their effectiveness. The percentage of effective nodules (NE) per plant was calculated based on the presence of brownish or pinkish pigmentation inside nodules. Thereafter, nodules were oven dried at 65°C for four days (Gwata ***et al***., 2004), and weighed to determine the total nodule dry weigh (NDW) per plant.

### DNA extraction

Young trifoliate leaves were collected from F2 plants, dried in Falcon tubes containing silica gel, and DNA was extracted using the cetyl trimethylammonium bromide (CTAB) method according to Kim ***et al***. (2012) with slight modifications. DNA pellet was re-suspended in 400 µL of 0.1× TE buffer, then 4 µL of RNAse-A was added to remove RNA. To ensure purity of DNA, further cleaning was done adding 40 µL Chloroform: Isoamyl alcohol (24:1) and mixing all thoroughly. Each Eppendorf tube was spun for 10 min. Then, fixed volume of 30 µL was picked from each sample and added equal volume of cold Isopropanol, then spun to pellet and washed in 70% ethanol, and air dried. Later, DNA was shipped to the National Soybean Research Laboratory (NSRL), University of Illinois, Urbana-Champaign, United States.

### Library preparation, genotyping and sequencing

GBS libraries are pools of short DNA fragments, obtained by the means of restriction enzymes, and ligated with barcoded adapters.

The work described here was carried out at the National Soybean Research Laboratory (NSRL) University of Illinois, Urbana-Champaign, United States, where all DNA was quantified using the intercalating dye (PicoGreen®; Invitrogen, Carlsbad, CA) and diluted to 25 ng/µL. A 96-plex GBS library comprising 92 soybean DNA samples, duplicate of parental lines and two negative (no DNA) controls were prepared as described in Elshire ***et al***. (2011). Five microliter (5µl) of each DNA sample was pipetted into wells of ready-to-use plate with A1 barcoded adapters at the bottom of each well, and 20µl restriction-ligation master mix was added to each well. The master mix contained two restriction enzymes; a rare cutter ***Hind***III/MseI, and a common cutter ***Hind***III***/Bfa***I, and a single restriction-ligation reaction was performed.

Samples were incubated and aliquot (50 µl) of each sample was pooled and cleaned up using 100ul AMPure beads. Pools were PCR amplified using 10uM Illumina primer pair (F+R) and 2X Phusion Master Mix (NEB #M0531), temperature cycling consisted of 98°C for 30s, followed by 18 cycles (98°C for 10s, 68°C for 30s, 72°C for 30s), and thereafter 72°C for 5min, and 4° C.

PCR products were run on agarose gel to verify smear of sheared DNA fragments, and a second cleanup was done using 100ul AMPure beads. DNA7500 chip was used for Agilent Bio Analyzer in order to determine average size (bp) and concentration (ng/µl) of fragments in the libraries.

Libraries were quantitated by qPCR (quantitative PCR) and sequenced on one lane for 101 cycles from one end of the fragments on a HiSeq2500, SBS sequencing kit version 4.

### Data processing: SNPS calling, alignment and SNPS production

Fastq.gz files were generated from the sequencing machine and de-multiplexed with the bcl2fastq v2.17.1.14 Conversion Software (Illumina). The sequencing result downloaded on the SFTP server (ftp://bdiers@ftp.biotec.illinois.edu) consisted in Fastq.gz files with over 267 millions of Reads (Reads are 100 nucleotides in length). Fatsq. gz files were processed and SNPs were called through the GBS analysis pipeline 2 as implemented in TASSEL 5.0 with modification of default settings; http://www.maizegenetics.net/tassel/docs/TasselPipelineGBS.pdf (Glaubitz ***et al***., 2014).

Briefly: i) tags were generated from fastq.gz files with the GBSSeqToTagDBPlugin (parameters: −KmerLength 80, -minKmerL 20, −c 10); ii) tags were extracted from database for alignment to reference genome with the TagExportToFastqPlugin (parameters: -c 1); iii) tags were aligned to the reference genome; http://soybase.org/cmap/cgibin/cmap/viewer?ref_map_set_aid=GmComposite2003;ref_map_aids=GmComposite2003_D1a;comparative_maps=1%3Dmap_aid%3DGmConsensus40_D1a;data_source=sbt_cmap. Bowtie2 version 2.2.8 was used for alignment; iv) tags were stored to physical map position in the database using the SAMToGBSdbPlugin, default parameters were used for that. v) SNPs were identified from the aligned tags with the DiscoverySNPCallerPluginV2 (parameters: mnLCov 0.05, mnMAF 0.01), vi) all discovered SNPs were scored for various parameters, including coverage, depth and genotypic statistics, using the SNPQualityProfilerPlugin, vii) based on the estimated parameters, quality scores were calculated using an R script, the inbreeding coefficient (minF), and the minimum minor allele frequency (minMAF) were both set to 0.05, viii) database was updated with quality scores using the UpdateSNPPositionQualityPlugin, ix) data from fastq and key files were converted and a VCF genotype file was generated (parameters: – kmerLength 80, and minPosQS 3). This generated VCF file contained the SNPs which need to be cleaned through further processing steps, as there are always a high number of missing SNPs, and some SNPs might have been affected by segregation distortion. Beagle version 4.1 (Browning and Browning, 2016) was used to impute missing SNPs, and ABH genotype file was generated and exported using the graphical interface of Tassel 5.0. (Bradbury ***et al***., 2007). Thereafter, a linkage map was generated using two R packages “ASMap” and “R-qtl”. Before building the linkage map, SNPs were filtered and cleaned up by removing low quality SNPs based on criteria such as “less than 10% missing data”, “duplicated SNPs”, and “SNPs with linkage distortion”.

### Data analysis

Phenotypic data were checked for normality, and data on nodules’ number (not normally distributed) were transformed using Box cox transformation (SAS Institute Inc. 2011). Box cox transformation tool suggested “square root” transformation as the most adequate. No transformation was more adequate than raw data for the other phenotypic traits (NE, NFW, NDW).

R-qtl (http://www.rqtl.org) was used to identify significant regions that are associated to promiscuous nodulation traits. After phenotype data was added to linkage map file, the genome was scanned via the “scanone” function of Haley-Knott (hk) regression, using the argument method="hk". Then, multiple imputation of the genome was performed using the function “sim. geno” with method="imp" and n. draws=64 (64 imputations). Thereafter, permutation test was performed to get a genome-wide significance threshold and p-values. Argument for that was method="hk" with n. perm= 1000 (1000 permutations). Two significance levels (alpha=0.2 and alpha=0.05) were set as suggested in the Rqtl manuel, the location of the QTL was estimated using the Haley-Knott (hk) regression method with both “lodint” and its alternative “bayesint” functions. Later, the closest flanking markers were identified, according to both intervals calculated in the former step. Lastly, the effects of the QTLs were estimated by plotting the average phenotype for each genotype group, the argument plotPXG was used for that purpose.

## RESULTS

Initially, a genotype file with over 65,000 SNPs was generated. After filtering low quality SNPs, a map with 29,620 SNPs covering all the twenty linkage groups was generated. Permutation tests generated various Lod score values for each trait. These ranged from 4.94 to 5.22 (Figure 1 and Table 1).

**TABLE 1:** Qtls position, and closest flanking markers for promiscuous nodulation traits

| Trait | Locus | Position | Threshold<br>(20%) | Lod<br>scores | P-values<br>($\alpha=0.2$) | Closest flanking markers |
| --- | --- | --- | --- | --- | --- | --- |
| NN | c10.loc3549 | 3549 | 4.99 | 5.05 | 0.181 | S10_33852650 and S10_29297382 |
|  | S13_11994810 | 7861 | 4.94 | 5.05 | 0.166 | S13_11994819 and S13_11994800 |
| NE | S13_11994810 | 7861 | 5.09 | 5.44 | 0.118 | S13_11994819 and S13_15403889 |
| NFW | c10.loc3545 | 3545 | 5.22 | 5.66 | 0.103 | S10_33852650 and S10_37822769 |
| NDW | c10.loc3545 | 3545 | 5.14 | 5.15 | 0.197 | S10_33852650 and S10_9412367 |

*NN= nodules' number, NE=percentage of effective nodules, NFW=fresh weight of nodules, and NDW=dry weight of nodules*

**FIGURE 1:**
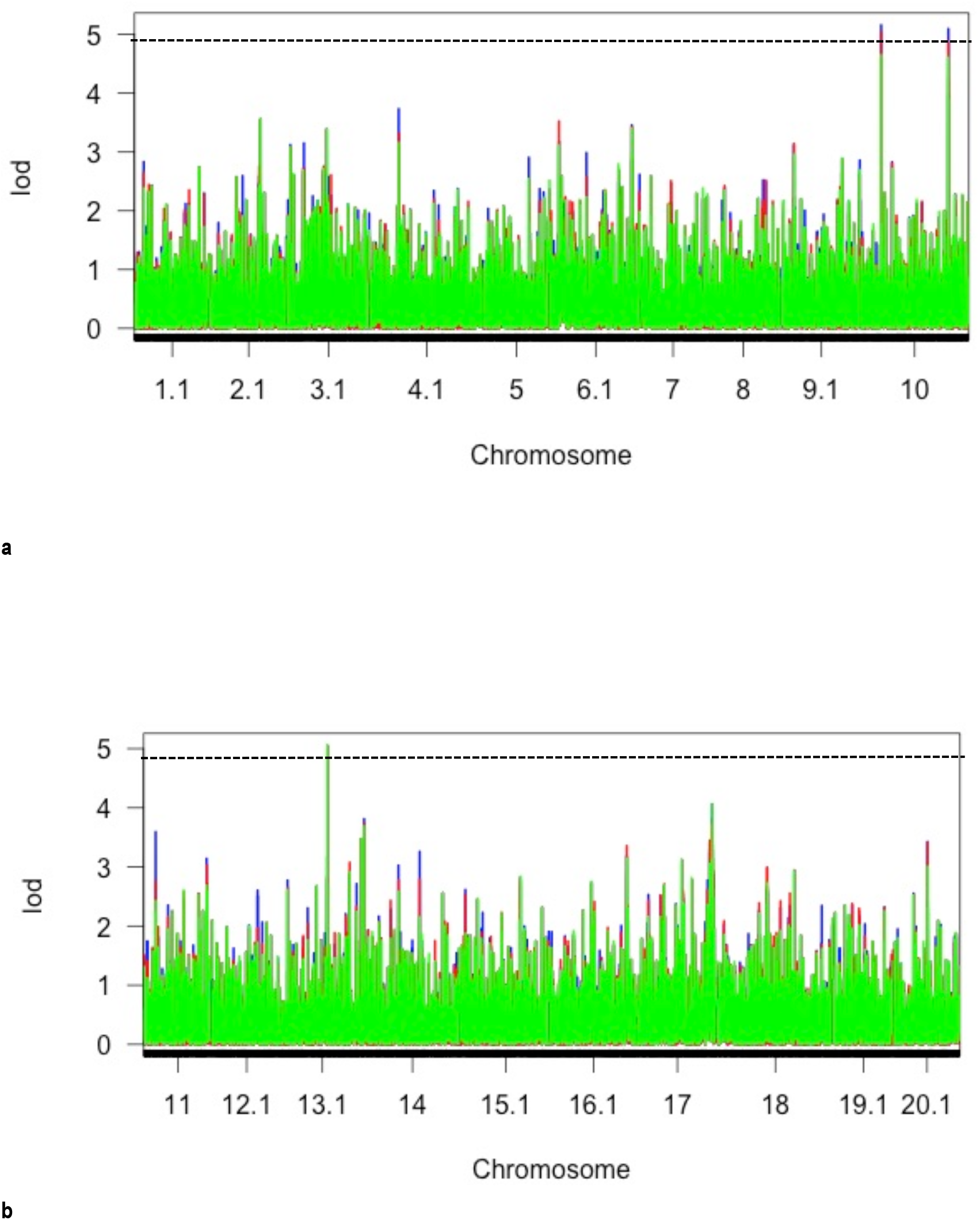
Curves showing Lod scores versus chromosomes: (a) chromosomes from 1 to 10 (b) chromosomes from 11 to 20

At least one significant QTL region was identified for all measured trait (Table 1). For the number of nodules (NN), two significant QTLs were observed: one located at the c10.loc3549 on the chromosome 10 (Lod score=5.05, and P-value=0.181). The marker S10_9400998 was the closest to that region, with S10_33852650 and S10_29297382 as flanking markers. The second QTL was located at the marker S13_11994810 on chromosome 13 (Lod score=5.05, and P-value=0.166), the flanking markers to that were S13_11994819 and S13_11994800.

For the percentage of effective nodules (NE), one QTL was found at S13_11994810 on chromosome 13 (Lod score=5.44, and P-value=0.118), with S13_11994819 and S13_15403889 as flanking markers.

The fresh weight of nodules (NFW) showed one significant QTL on chromosome 10 at the loci c10.loc3545 (Lod score=5.66, and P-value=0.103), the flanking markers were S10_33852650 and S10_37822769.

The dry weight of nodules also exhibited one significant QTL on chromosome 10 at the loci c10.loc3545 (Lod score=5.15, and P-value=0.197), with S10_33852650 and S10_9412367 as flanking markers.

As for the QTL effects, Figure 2a showed higher mean performance for the progenies compared to the parents’ mean, suggesting over dominance QTL effect on nodules’ number on chromosome 10. On the chromosome 13 (Figure 2b) the mean performance for progenies were lower compared to both parents. This suggest non-additive QTL effect on number of nodules. For nodules’ effectiveness (NE), only one QTL was found, and this was on chromosome 13. Similar to the situation observed in NN on that chromosome, progenies mean performance were lower than both parents’ (Figure 2c). As for the fresh weight of nodules (NFW), one QTL was found on chromosome 10, progenies had higher mean performance than both parents (Figure 2d), this situation is similar to the one observed in NN on the same chromosome 10, suggesting an over dominance QTL effect on NFW. The same apply to dry weight of nodules (NDW) (Figure 2e), with one QTL on chromosome 10, where progenies’ mean performance were higher than both parents’.

**FIGURE 2:**
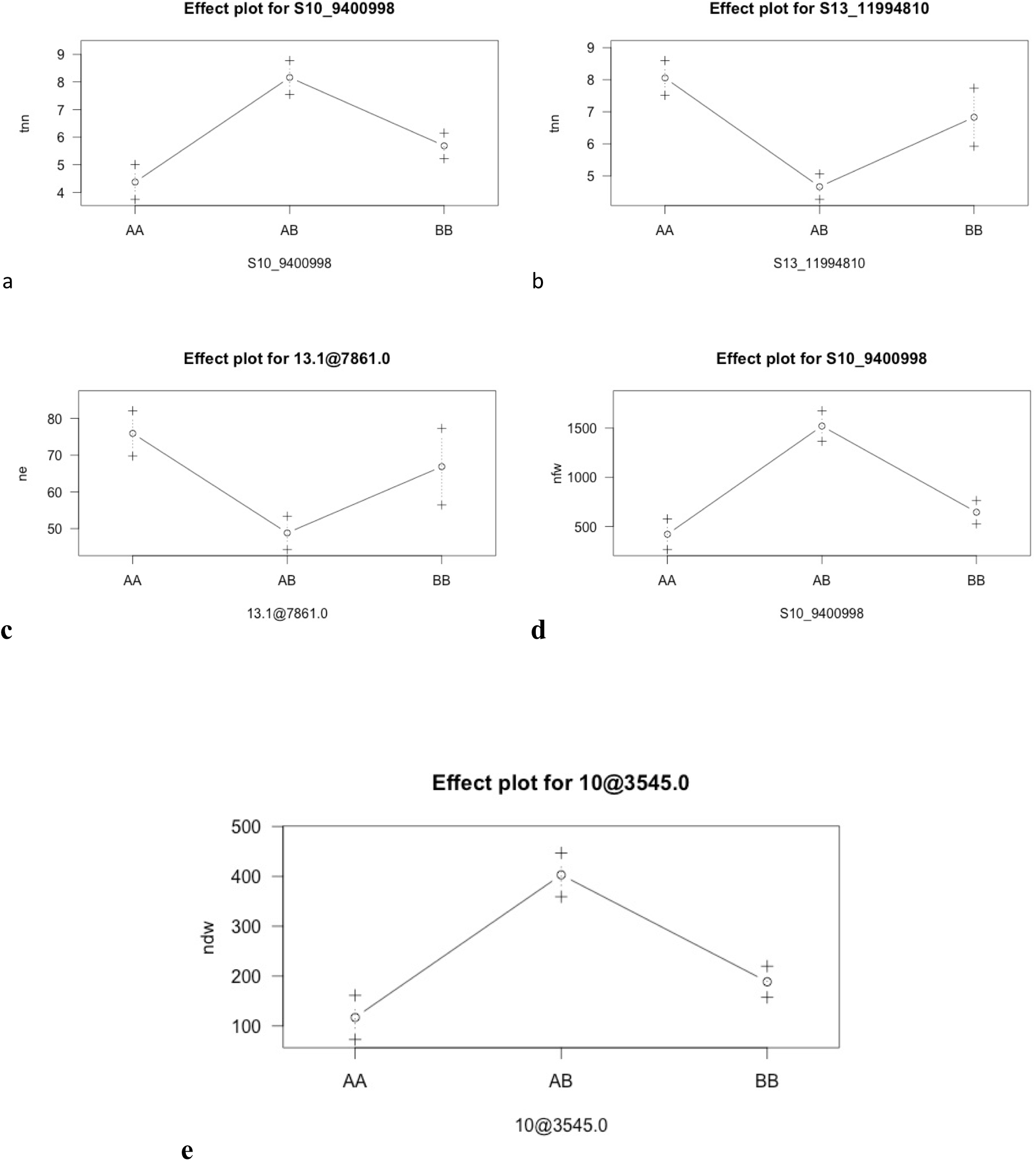
QTLs effects for nodule traits on chromosome 10 and 13

## DISCUSSION

The QTL identified on chromosome 10 for NN was the same as the one of NFW and NDW. Even though the closest flanking markers were not exactly the same, they all shared one closest flanking marker, namely S10_33852650. This is an indication that these three traits are under the control of the same genome region, suggesting a possibility of pleiotropic genes in that region. The same was observed on chromosome 13 for NN and NE, where the marker S13_11994810 was significant for both traits at position 7861. Although the flanking markers showed larger interval for NE, both traits shared the flanking marker S13_11994819. As observed previously, this suggests the probable existence of pleiotropic genes in that region, which control nodule number and effective nodules. The genotypes wondersoya and CG 2043 used in this study revealed the existence of QTLs associated with promiscuous nodulation in soybean, these findings open doors to the possibility to carry out marker assisted breeding. However, the over dominance effect of QTL consistently observed on chromosome 10 is an indication of interactions of several alleles at this region. As described by East (1908) and Shull (1908), over dominance is a result of cumulative action of divergent alleles or stimulation of divergent alleles in heterozygotes. This hypothesis was supported by Hull (1945). East (1936) emphasized further that a combination of higher number of divergent alleles exhibit higher heterosis than less divergent combinations. Hence, this study enabled understand that in promiscuous nodulation, heterozygotes can have advantage to the survival of many alleles that are recessive and harmful in homozygotes. The detection of those QTLs that enable the above mentioned hypotheses is very insightful, as it may inform on the possibility applying marker assisted selection on promiscuity character in soybean breeding, also this can serve as starting point for gene cloning.

The fact that no significant QTL was observed with p-value =0.05 can be explained by the low population size (92 individuals). For instance, Semagn ***et al.*** (2006) reported that a total of 200 individuals is required to construct a reasonably accurate linkage map. Also some of the SNPs closer to the interesting regions could been filtered out during data processing, leading to reduction in accuracy. It is therefore recommended that SSR markers that are polymorphic in those regions be tested to confirm these observations. Moreover, it would be useful to perform such QTL identification study on a larger F2 population or RILs population, as this would yield more QTLs with more accuracy. It appears that a set of genes located in those genomic regions on the chromosomes 10 and 13 control the nodulation traits, this suggest the hypothesis of pleiotropic genes involved in the expression of the nodulation traits. It is recommended that those genes be cloned for proper understanding of the promiscuity character in soybean.

## CONCLUSION

There are quantitative traits loci associated with promiscuous nodulation in soybean. The present study identified two loci, one on chromosome 10 (c10.loc3545) and one on chromosome 13 (S13_11994810). All nodulation traits were associated with one or both QTLs. Number of nodules (NN) was associated with both QTLs. NN shared one QTL (c10.loc3545) with NFW and NDW, and the QTL (S13_11994810) with NE. This allows inferring on the possibility of applying maker assisted selection and search for pleiotropic genes in those genomic regions. Also, the findings of this study can serve as starting point for gene cloning. Therefore, it is recommended that investigations be pursued on that matter, to identify and clone those genes to enable boosting breeding activities for the development of promiscuous soybean cultivars.

## ACKNOWLEDGEMENT

Eric Agoyi is a Fellow of the Norman E. Borlaug Leadership Enhancement in Agriculture Program funded by USAID. Support for this research was provided in part by the Borlaug Leadership Enhancement in Agriculture Program (Borlaug Leap) through a grant to the university of California Davis by the United States Agency for International Development. The opinions expressed herein are those of the authors and do not necessarily reflect the views of USAID. Acknowledge goes to Prof. Diers’ Lab in the National Soybean Research Laboratory (NSRL) at the University of Illinois, Urbana-Champaign, United State of America for providing lab facilities and technical support to the im[plementation of this research. Wlso we acknowledge support received from NaCCRI and CIAT-Uganda for their support during DNA extraction and processing for shipment.

